# Compaction dynamics during progenitor cell self-assembly reveal granular mechanics

**DOI:** 10.1101/699447

**Authors:** Bart Smeets, Jiří Pešek, Thomas Deckers, Gabriella Nilsson Hall, Maxim Cuvelier, Steven Ongenae, Veerle Bloemen, Frank P Luyten, Ioannis Papantoniou, Herman Ramon

## Abstract

We study the self-assembly dynamics of human progenitor cells in agarose micro-wells that are used for production of chondrogenic organoids. Using image analysis on time-lapse microscopy, we estimate the aggregate area in function of time for a large number of aggregates. In control conditions, the aggregate radius follows an exponential relaxation that is consistent with the dewetting dynamics of a liquid film. Introducing Y-27632 Rho kinase inhibitor, the compatibility with the liquid model is lost, and slowed down relaxation dynamics are observed. We demonstrate that these aggregates behave as granular piles undergoing compaction, with a density relaxation that follows a stretched exponential. Using simulations with an individual cell-based model, we construct a phase diagram of cell aggregates that suggests that the aggregate in presence of Rho kinase inhibitor approaches the glass transition.

## Introduction

Recent years have witnessed revolutionary advancements in the field of 3D stem cell research through the development of strategies that exploit initial cellular aggregation as a means to obtain functional tissue structures termed “organoids”. Organoid technologies rely on the exploitation of the intrinsic capacity of stem cells to self-assemble into aggregates and condense. They can be used for engineering tissue implants [1] and for disease modeling [2]. Using such formats, the mimicry of developmental cascades through consecutive self-patterning processes has been carried out using adult stem cells or pluripotent stem cells, resulting in organoids of various tissue types such as intestine, liver, pancreas, kidney, prostate, lung, optic cup, and brain [3]. In micro-aggregation culture formats, the number of cells in each aggregate can be accurately controlled [4]. This facilitates the detailed investigation of cell assemblies, and allows the high-throughput generation of configurations that compare in dimension to in vivo developmental condensation events [5, 6]. The ability to understand the compaction process that initiates cellular self-assembly is crucial, as it has been linked to stem-cell lineage specification and commitment [4]. Aggregate compaction is a function of multiple cellular processes such as cell movement and shape control [7], cell-cell communication [8], cell-substrate interactions [9], tissue biomechanics and mechanotransduction [10].

The compaction that takes place in cell aggregates has been compared to the classical dewetting transition. Analogous to a liquid droplet, the balance of cell-cell and cell-substrate adhesion energy governs whether an aggregate will spread or form a 3D droplet [11]. Using this analogy, the spreading radius and contact angle have been successfully related to cohesive and adhesive properties of epithelia [12]. By studying their deformation dynamics, the relative viscosity of spreading cell aggregates could be deduced [11]. On adhesive substrates, it has been shown that shape fluctuations caused by active tractions may induce morphological instabilities that introduce a colony size dependency to the wetting transition [13]. On soft and/or weakly adhesive substrates, cells are unable to generate significant substrate traction forces. As a result, the dewetted state is always favored in these conditions [14, 15]. In this paper, we study the compaction dynamics of aggregates from human periosteal derived cells (hPDC), a population of human progenitor cells, in agarose micro-wells, a soft and weakly adhesive substrate that is commonly used for the production of self-assembled micro-aggregates.

## Results

### Aggregate formation dynamics of progenitor cells are consistent with liquid dewetting

Edwards et al. [16] analyzed the dewetting of a liquid film into a single drop. For a large thin film, they identified two dynamic regimes: First, the radius decreases linearly in time, accompanied by an annular rim close to the contact line. Next, the rim merges, and the film relaxes exponentially to a spherical cap shape. We seeded human progenitor cells at sufficient density on a non-adhesive agarose substrate. Within a few hours, the film of cells starts to contract to a condensed clump, Fig. 1(a). Although this contraction occurs with a peripheral thickened rim, reminiscent of the linear regime of the dewetting liquid film, this experiment is not ideal for a quantitative comparison with liquid theory, as the time-scale of the contraction is too large to rule out influential biological processes such as cell division, cell death or phenotypic changes. Moreover, the granular nature of the film makes it hard to ensure a uniform cell distribution, leading to gaps and a fractal structure near the edge of the film.

**Figure 1:**
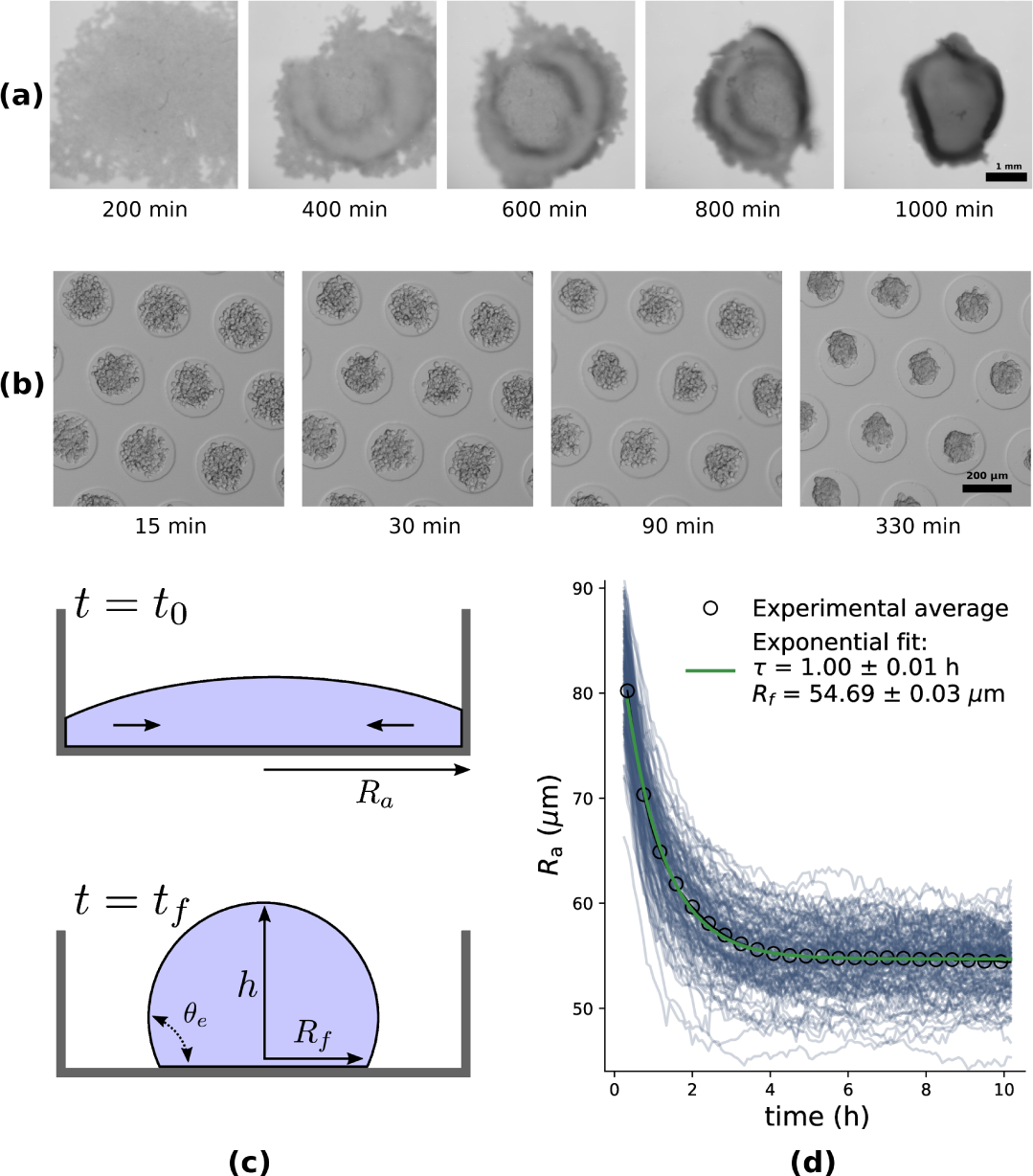
Dewetting dynamics of condensing aggregates. **(a)**: Time-lapse microscopy of a large film of progenitor cells that is dewetting from an agarose substrate. At intermediate times, a characteristic thickened ‘rim’ can be observed near the periphery of the cell aggregate. **(b)**: Time-lapse microscopy images of aggregate formation of progenitor cells in agarose microwells at logarithmically spaced times. **(c)**: Illustration of the dewetting of a liquid droplet with radius *R*_*a*_(*t*). At *t* = *t*_*f*_, the droplet shape is equilibrated with final radius *R*_*f*_ and contact angle *θ*_*e*_. **(d)**: Equivalent aggregate radius over time, sampled every 5 minutes during early aggregate compaction (10 h). Each blue line represents a single analyzed aggregate out of 195 analyses. The black circles indicate the binned average of these, and the full green line shows the exponential fit.

Fig. 1(b) shows an alternative experimental setup that is better suitable for a quantitative comparison. We seed hPDCs at a controlled cell density in small agarose micro-wells with a diameter of around 200 µm. Initially, the cells are nearly uniformly distributed in each well. With time, they contract into small circular aggregates. This phenomenon is similar to the final relaxation phase of a dewetting liquid film, as illustrated in Fig. 1(c). For this regime, Edwards et al. [16] found an analytical expression, relating the exponential relaxation time *τ* to the viscosity *η*_*a*_, the surface tension *γ*_*a*_ and the volume *V*_*a*_ of the aggregate, namely

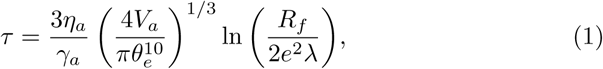

where *R*_*f*_ is the final contact radius of the dewetted spherical cap, *θ*_*e*_ is the apparent liquid-solid contact angle – see Fig. 1(c) – and *λ* is the (Navier) slip length of the viscous fluid. At long timescales (minutes to hours) multicellular masses behave as highly viscous fluids with surface tension enacted by intercellular adhesion energy [17]. When applying (1) to micro-aggregates, *η*_*a*_ represents the bulk apparent viscosity and *γ*_*a*_ the net surface tension.

We analyzed the area over time of 195 independent microwells containing 200 hPDCs each. The average temporal profile of the associated radii showcases a remarkably clear exponential relaxation – Fig. 1(d). By fitting an exponential decay to this curve we estimate *τ* = 1.00 ± 0.01 h and *R*_*f*_ = 54.69 ±0.03 µm. The total volume can be approximated based on the average number of cells per well and the measured sizes of individual cells, see SI Fig. 4, assuming that cells maintain their initial volume. For 200 cells, we thus compute *V*_*a*_ ≈ 6.1×10^5^ µm^3^. Given the slip length *λ* and the wetting angle *θ*_*e*_, the ratio *γ*_*a*_*/η*_*a*_ can be obtained from (1). The slip length is a microscopic property that cannot be rigorously defined nor experimentally measured for a coarse packing of cells, with complex non-isotropic material properties near the interface. However, it only enters (1) through a weak logarithmic dependency. Hence, an order-of-magnitude estimation suffices. At the cell-substrate interface we can write *λ* = *v*_wall_ *η*_wall_*/τ*_wall_, with *v*_wall_ the local slip velocity, *η*_wall_ the apparent microscopic viscosity and *τ*_wall_ the shear stress. Magnetic bead rheometry experiments on adhering fibroblasts in [18], using a sliding velocity *v*_wall_ ≈1.9 µm/min and an applied shear stress of *τ*_wall_ ≈ 300 Pa found an apparent local viscosity of *η*_wall_ ≈ 2 kPa*·*s. This results in *λ* ≈ 215 nm, roughly the thickness of the acto-myosin cortex [19, 20]. We estimate the apparent wetting angle *θ*_*e*_ by assuming that the final aggregate is a spherical cap with volume *V*_*a*_ and with radius *R*_*f*_, hence *θ*_*e*_ ≈ 108°, corresponding to an aggregate aspect ratio of 0.69. Then, we obtain from (1) *γ*_*a*_*/η*_*a*_ ≈ 1.58 µm/min, the characteristic relaxation speed of the tissue fluid. Guevorkian et al. [17] used pipette aspiration experiments on small spheroids of S180 cells to determine their apparent viscosity and surface tension. They found *γ*_*a*_ = 6 mN/m and *η*_*a*_ = 1.9×10^5^ Pa*·*s, hence a relaxation speed of *γ*_*a*_*/η*_*a*_ = 1.89 µm/min, a value well comparable in magnitude to our estimate using (1). Consequently, this analysis suggests that the *average* aggregate formation dynamics are consistent with the dewetting of a liquid film that is compatible with rheological tissue properties.

### Rho kinase inhibitor treatment induces granular compaction dynamics

We repeated the micro-aggregation experiment, but for aggregates that have been treated with Rho kinase inhibitor (further denoted as Rocki) Y-27632 at concentrations of 10 and 20 µM in the aggregation medium (see SI Fig. 1 and 2). Confocal microscopy shows that the treatment with Rocki leads to more granular aggregate morphology, with more rounded cells and a more irregular aggregate shape – Fig. 2. Again, we analyzed the temporal dynamics of the corresponding aggregate radii, shown in Fig. 3(a). Surprisingly, the close correspondence of the exponential fit is lost for the aggregates treated with Rocki – Fig. 3(a).

**Figure 2:**
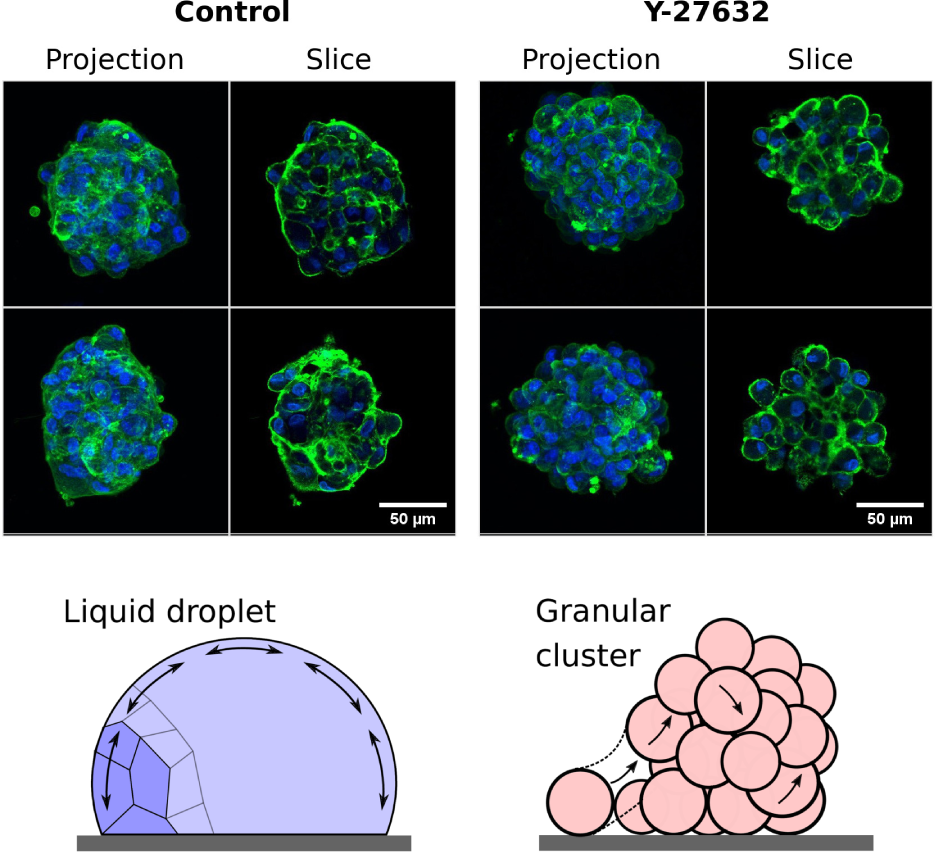
Rho kinase inhibitor affects aggregate morphology. Top: Representative confocal microscopy images of aggregates 10 h after seeding, in DMEMc medium with 0 µM and 10 µM Y-27632 Rho kinase inhibitor, stained for F-actin (phalloidin, green) and nucleus (DAPI, blue). For each aggregate, both the maximum intensity projection (projection) and a z-section (slice) are shown. Bottom: Illustration of corresponding limiting phases of cell aggregates: In a granular cluster, the displacement of individual cells is decoupled, causing active jamming behavior and slow power law-like scaling of the aggregate radius. In the fluid phase, microscopic deviations are quickly equilibrated, and a globally established tension governs the exponential aggregate relaxation.

**Figure 3:**
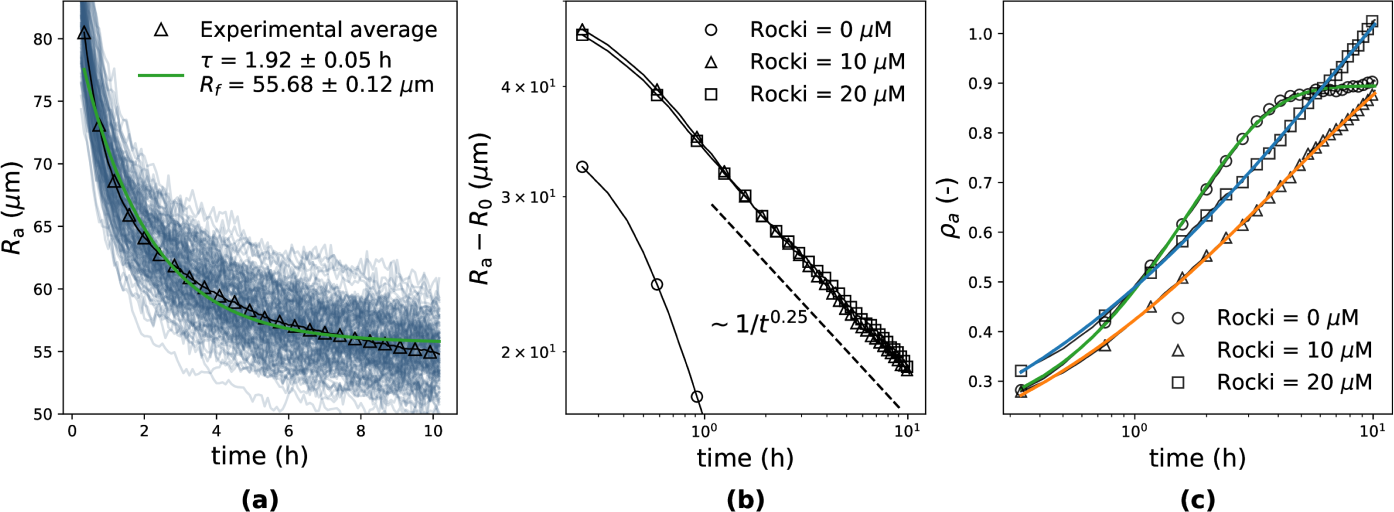
Rocki affects aggregate compaction dynamics. **(a)**: Aggregate radius in function of time for 10 µM Rocki. Each blue line is a single aggregate of ± 200 cells during 10 h. Triangles indicate the average aggregate radius at each time point. The full green line shows the exponential fit. **(b)**: Double-logarithmic plot of the shifted aggregate radius *R*_*a*_ − *R*_0_ as a function of time, comparing Rocki = 0 µM (circles), Rocki = 10 µM (triangles) and Rocki = 20 µM (squares). For each condition, *R*_0_ was determined from the power law fit *R*_*a*_(*t*) − *R*_0_ = *b t*^*−β*^*P*. **(c)**: Estimated apparent packing density *ρ*_*a*_ as a function of time, varying Rocki, including fits of the KWW model. The values and standard deviations of the estimated parameters are listed in Table 1.

**Figure 4:**
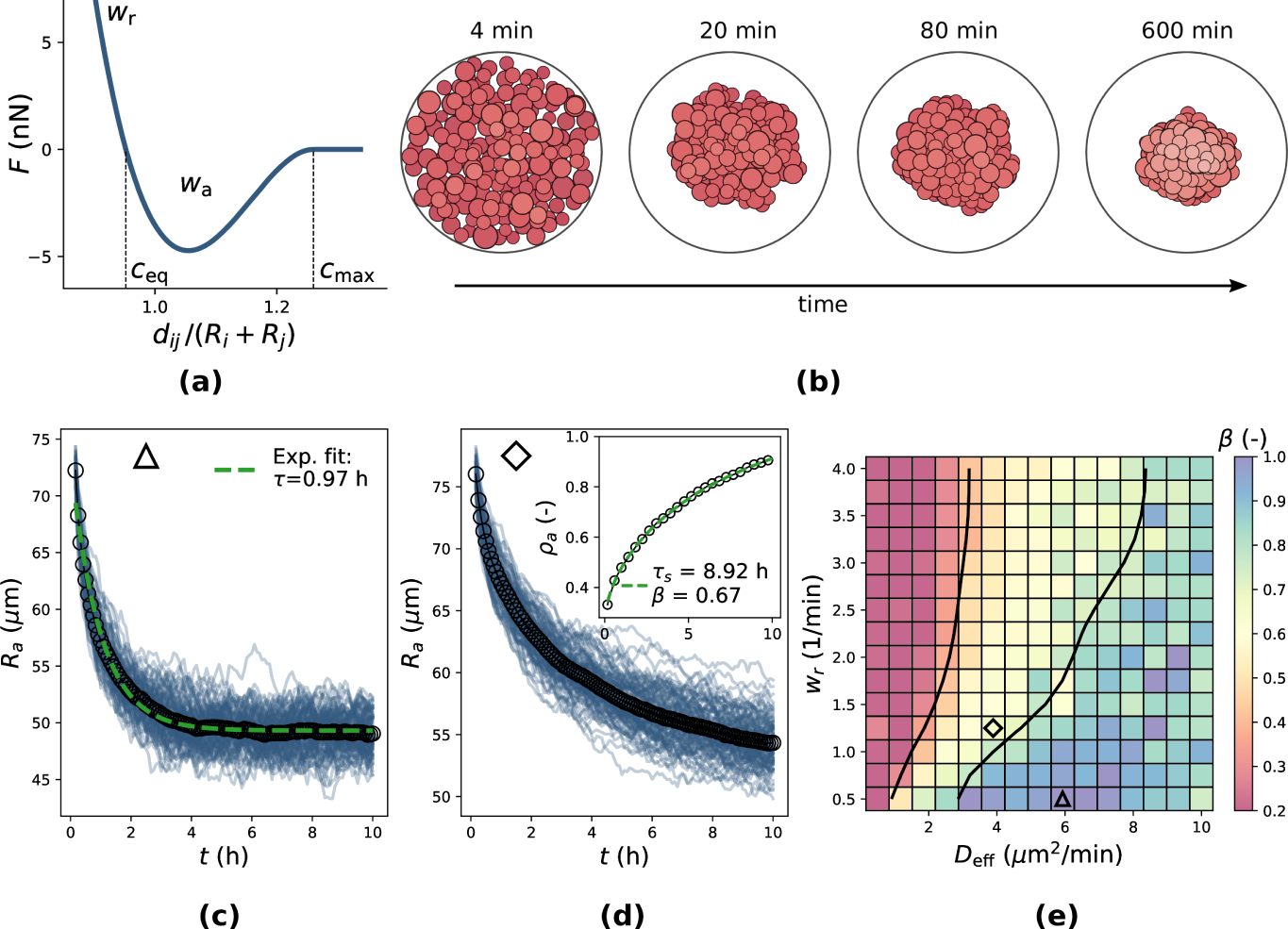
Individual cell-based simulation of aggregate formation. **(a)**: Illustration of ‘soft’ inter-cellular force model based on adhesive tension *w*_*a*_ and stiffness *w*_*r*_ (cortical tension). **(b)**: Visualization of simulated aggregate compaction process, starting from deposited cells in the micro-well (see SI Fig. 3) at different, logarithmically spaced time points. The microwell has a diameter of 200 µm. **(c)** and **(d)** Change of radius as a function of time for simulated aggregates in both liquid **(c)** and supercooled liquid **(d)** regimes. **(e)**: Map of estimated exponent of (2) for varying stiffness *w*_*r*_ and cell activity *D*_eff_. Blue values indicate liquid-like behavior while red values indicate glassy or jammed states. The black lines are the isocontours at *β* = 0.3 and *β* = 0.75, roughly indicating the respective transitions between glass, supercooled liquid and liquid. The triangle and diamond symbols indicate the locations in the map from where resp. **(c)** and **(d)** were sampled from. Fig. **(c-e)** were simulated at *w*_*a*_ = 4 min^−1^, *D*_*r*_ = 0.5 min^−1^. For each parameter combination, 100 independent simulations were performed, of which the average was used to fit (2) and estimate *β*.

Instead, power law relaxation of the aggregate radius is observed with *R*_*a*_(*t*) − *R*_0_ ∼ *t*^−0.25^, Fig. 3(b). This slow relaxation is reminiscent of aging in colloidal glasses and gels [21], and the compaction of jammed granular material [22, 23], see Fig. 2. The relaxation dynamics during compaction of granular media have been well studied [23]. For instance, the density relaxation of glass beads in a vibrating vessel follows the Kohlrausch-Williams-Watts (KWW) law, a stretched exponential, of the apparent packing density *ρ*_*a*_ [24]:

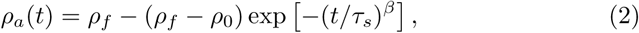

where *ρ*_*f*_ and *ρ*_0_ are respectively the steady state and initial packing fraction, and *τ*_*s*_ and *β* are the characteristic timescale and relaxation exponent. Given the aggregate radius *R*_*a*_ and the known average volume of cells in each aggregate, *V*_*a*_, we calculated the apparent packing density 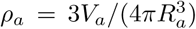. Fig. 3(c) shows the time evolution of *ρ*_*a*_ for varying Rocki with stretched exponential fits. The fitted parameters are listed in Table 1. The experimental data is in excellent agreement with (2) for all three Rocki conditions. The exponent *β* is used in the study of spin glasses, clusters of soft spheres and polymers to delimit the glass transition [25, 26]. At high temperature, *β* ≈ 1, and the material behaves as a granular fluid with exponential relaxation of *ρ*_*a*_. As the liquid becomes supercooled with decreasing temperature, *β* starts to decrease until it reaches the glass transition [26]. The value of *β* where this occurs depends on the model system (and finite system size), but is generally between *β* = 1*/*3 and *β* = 0.6 [26]. We find that *β >* 1.0 for Rocki = 0 µM, consistent with the suitability of a fluid model (see further). For Rocki = 10 µM and Rocki = 20 µM we find values of *β* ≈ 0.5, Table 1.

**Table 1:**
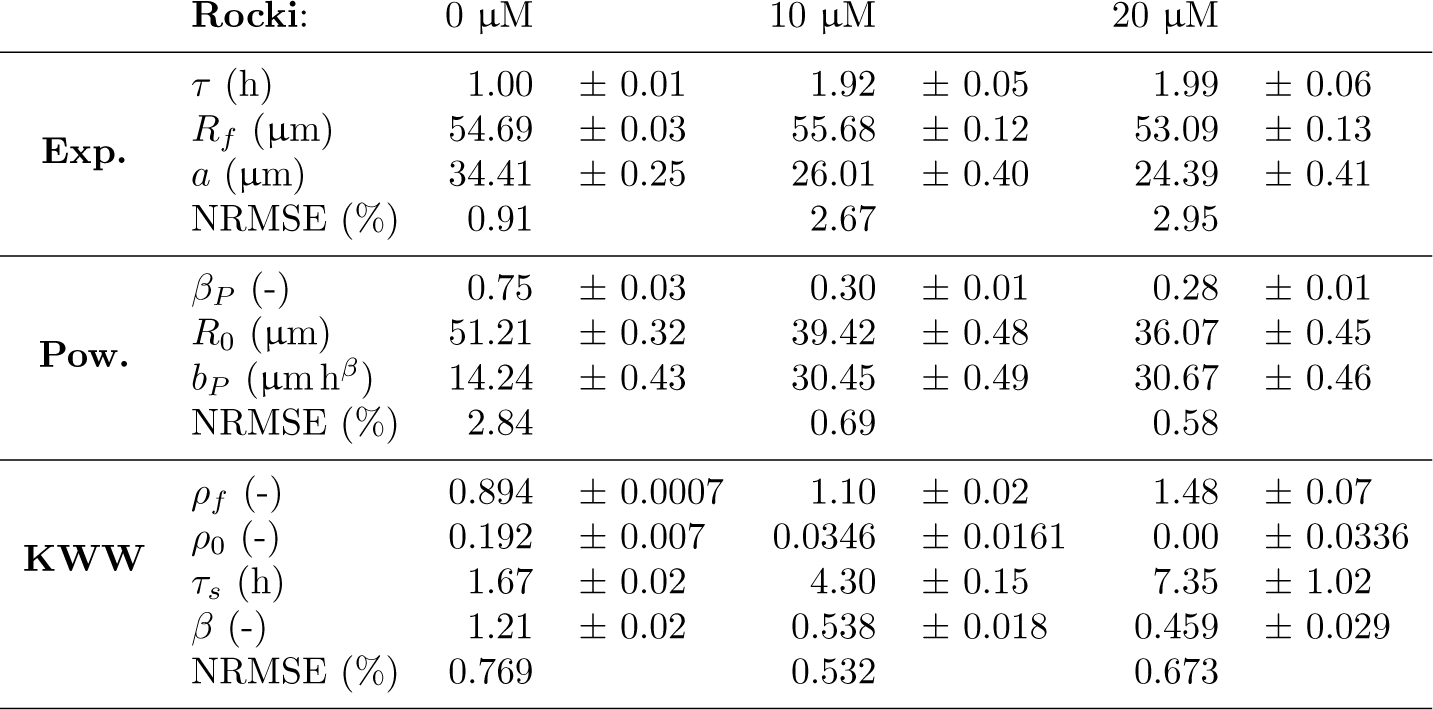
Parameters for exponential fit (Exp.) *R*_*a*_(*t*) *- R*_*f*_ = *a* exp (*-t/τ*), power law fit (Pow.) *R*_*a*_(*t*) *- R*_0_ = *b t*^*-β*^*P* and KWW fit, (2), on aggregate radius vs. time varying Rocki. Standard errors are indicated for each parameter estimate, obtained from the covariance matrix of the non-linear least squares fit. NRMSE is the normalized root mean square error.

| | Rocki: | 0 $\mu\text{M}$ | | 10 $\mu\text{M}$ | | 20 $\mu\text{M}$ | |
| --- | --- | --- | --- | --- | --- | --- | --- |
| **Exp.** | $\tau$ (h) | 1.00 | $\pm 0.01$ | 1.92 | $\pm 0.05$ | 1.99 | $\pm 0.06$ |
| | $R_f$ ($\mu\text{m}$) | 54.69 | $\pm 0.03$ | 55.68 | $\pm 0.12$ | 53.09 | $\pm 0.13$ |
| | $a$ ($\mu\text{m}$) | 34.41 | $\pm 0.25$ | 26.01 | $\pm 0.40$ | 24.39 | $\pm 0.41$ |
|  | NRMSE (%) | 0.91 |  | 2.67 |  | 2.95 |  |
| **Pow.** | $\beta_P$ (-) | 0.75 | $\pm 0.03$ | 0.30 | $\pm 0.01$ | 0.28 | $\pm 0.01$ |
| | $R_0$ ($\mu\text{m}$) | 51.21 | $\pm 0.32$ | 39.42 | $\pm 0.48$ | 36.07 | $\pm 0.45$ |
| | $b_P$ ($\mu\text{m h}^\beta$) | 14.24 | $\pm 0.43$ | 30.45 | $\pm 0.49$ | 30.67 | $\pm 0.46$ |
|  | NRMSE (%) | 2.84 |  | 0.69 |  | 0.58 |  |
| **KWW** | $\rho_f$ (-) | 0.894 | $\pm 0.0007$ | 1.10 | $\pm 0.02$ | 1.48 | $\pm 0.07$ |
| | $\rho_0$ (-) | 0.192 | $\pm 0.007$ | 0.0346 | $\pm 0.0161$ | 0.00 | $\pm 0.0336$ |
| | $\tau_s$ (h) | 1.67 | $\pm 0.02$ | 4.30 | $\pm 0.15$ | 7.35 | $\pm 1.02$ |
| | $\beta$ (-) | 1.21 | $\pm 0.02$ | 0.538 | $\pm 0.018$ | 0.459 | $\pm 0.029$ |
|  | NRMSE (%) | 0.769 |  | 0.532 |  | 0.673 |  |

To investigate the transition in more detail, we created a minimal individual cell-based model (IBM) of the compaction process, based on mechanical forces exchanged between cells. The IBM is derived from the model introduced by Delile et al. [27], and is akin to Self-Propelled Particle (SPP) simulations [28, 29], with the important difference that active forces are conservative and exchanged between neighboring cells. A full description of the model can be found in the supplementary information. Briefly, it has the following attributes: Cells are represented by their central coordinates; symmetric cell-cell contact forces are computed for all edges of the Delaunay triangulation of these coordinates and based on a smooth continuous central potential parameterized by adhesion and cortical tension as illustrated in Fig. 4(a); cells carry a randomly diffusing direction vector representing their polarization; protrusive forces are computed between neighboring cells based on their polarization and an active traction parameter; finally, the system is evolved over time by integrating overdamped equations of motion. Crucially, and contrary to classical Active Brownian Particle (ABP) models with hard-sphere potentials, the (soft) interaction potential and the Delaunay-based contact resolution scheme were designed to simulate a granular fluid at confluent cell densities. After rescaling the model parameters to represent the experimentally accessible data, the IBM characterizes the aggregate compaction in four main mechanical properties: the repolarization frequency *D*_*r*_ (min^−1^), inversely proportional to the cell’s persistence, the active cell velocity *v*_*t*_ (µm/min) that causes mixing of the cells in the aggregate, and the repulsive and adhesive relaxation rates *w*_*r*_ and *w*_*a*_ (min^−1^) which parameterize the condensation rate of a doublet of adhering cells from the balance of cortical tension, adhesion and apparent cytoskeletal viscosity. Finally, using *v*_*t*_ and *D*_*r*_, we define 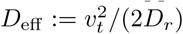 as a measure for active cell motility (see SI Fig. 6).

Fig. 4(b) visualizes a simulated aggregate compaction process. For high levels of cell activity *D*_eff_ and soft repulsion *w*_*r*_, we expect liquid properties, while for reduced activity and stiff repulsion, we expect glassy behavior. Indeed, simulations produce exponential relaxation dynamics of the aggregate radius at large *D*_eff_ and low *w*_*r*_ – Fig. 4(c). At lower *D*_eff_ and higher *w*_*r*_, the observed relaxation of the aggregate radius is not exponential – Fig. 4(d). However, all simulations agree well with the KWW law for granular compaction. This is further confirmed in Fig. 5 which shows the time evolution of *ρ*_*a*_ for varying *D*_eff_ (∼ temperature), illustrating the transition from a fluid to a colloidal glass. It should be noted that the distinction between a supercooled liquid and a glass is not strict but depends on the relevant timescale, in this case the observation time of the aggregate formation process [30]. Maximal compaction (or conversely, smallest aggregate size) is attained for intermediate values of *D*_eff_. Indeed, in the presence of too much agitation, the outward swim pressure established by active forces inflates the aggregate beyond the minimal size imposed by the interaction potential [31].

**Figure 5:**
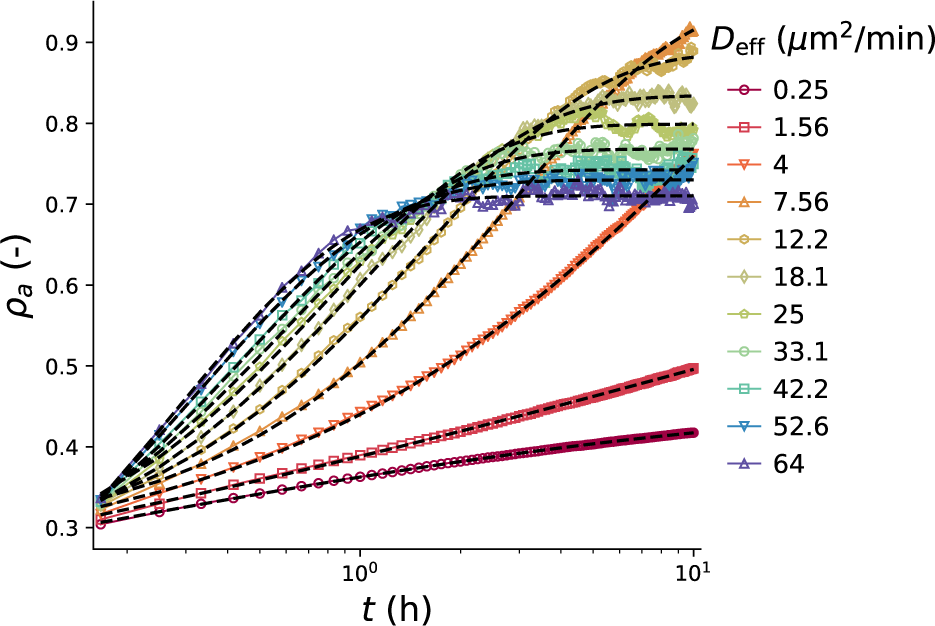
Packing density in simulated aggregates. Packing density *ρ*_*a*_ as a function of time, for simulated aggregates with varying effective cell diffusivity 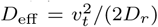. Each curve is obtained as the average over 150 independent simulations of a single cell aggregate. Dashed black lines indicate the fit using the stretched exponential (2). *D*_eff_ was modulated by varying *v*_*t*_, keeping *D*_*r*_ constant at 0.5 min^−1^. Furthermore, we set *w*_*a*_ = 4 min^−1^ and *w*_*r*_ = 2 min^−1^.

The estimated values of *β* using (2) for varying *D*_eff_ and *w*_*r*_ are shown in Fig. 4(e). This provides an approximate phase diagram of cell aggregates with the following regions: A fluid at large *D*_eff_ and low *w*_*r*_, where *β* ≈ 1, a supercooled fluid at intermediate *D*_eff_ and *w*_*r*_, wherein *β* decreases and finally a colloidal glass at low *D*_eff_ and large *w*_*r*_, with *β* ≈ 0.3. We have focused on the parameters *D*_eff_ and *w*_*r*_, due to their parallel in the glass transition with temperature and density [32]. The total cell-cell potential is also determined by the strength of adhesion *w*_*a*_. In the SI, Fig. 7, we conclude that *w*_*a*_, if sufficiently high, contributes to the material properties in a multiplicative manner, leading to a net potential strength 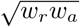.

A comparison of the experimental estimates of *β* with this diagram suggests that for the control condition at Rocki = 0 µM, compaction dynamics most resemble those of a granular liquid. Interestingly, in the experiment we estimated *β* = 1.21 (Table 1), while *β* does not exceed one for any parameter combination in our simulations. This yet unexplained super-exponential relaxation of the packing factor could be attributed to changes that occur at the cellular level, for example the maturation of cell-cell adhesion over time [33], or the gradual decrease of cell volume of the cells that are compressed in the aggregate [34]. For the experiments including Rocki, we estimated that *β* ≈ 0.5 (Table 1), which is consistent with a supercooled granular fluid that is approaching the glass transition – Fig. 4(e).

## Discussion

While the mechanical properties of biological cells and tissues are characterized by complex non-linear and time-dependent phenomena [35], simple models of viscous liquids with surface tension have been remarkably successful at describing their properties at intermediate to long timescales (minutes to hours) [36, 11]. In this work, we performed an analysis of micro-well aggregate formation from a population of human progenitor cells. Micro-wells are a commonly used culture platform for tissue engineering applications. The high throughput approach has made it possible to acquire a large number of samples (> 100) for each experiment, permitting us to distinguish and interpret subtle differences (see Fig. 1) between experimental conditions with unusually large discriminatory power.

We found that in control conditions, the dynamics of micro-aggregate formation are compatible with the model of a liquid with surface tension. However, when perturbing the system with Rho kinase inhibitor, this correspondence is lost. Instead, we observed slowed down relaxation dynamics characteristic of glassy systems. The liquid-glass and closely linked liquid-jamming transition have been extensively studied, both theoretically [37] and experimentally [38, 39]. Jamming has recently been studied in the context of biological tissues [40, 41, 42] to help elucidate phenomena such as collective migration [43], embryonic development [44] and cancer [45, 46]. These studies investigate the jamming phenomenon for confluent tissues that are simulated using some variant of vertex models.

This is the first work that investigates the liquid-glass transition in aggregates used to produce organoids for tissue engineering applications using adult human progenitor cells, an inherently complex cell population. Findings in this study are expected to be applicable also to other progenitor populations derived from the bone marrow and adipose tissue. We demonstrated that the apparent packing density follows a stretched exponential, in excellent agreement with the relaxation dynamics during compaction of a granular medium [23]. This agreement is found across three experimental conditions. This suggests that (early) cell aggregates can be described as agitated granular piles. This agitation is caused by cell activity, which regulates whether the pile transitions from a granular fluid over a supercooled fluid to a glass. By means of individual cell-based simulations of the aggregate formation, we confirmed that this transition is contingent on the level of cell activity and the repulsive strength of cell-cell interactions. This finding is in line with the phase diagram for active dense tissues proposed in [41], where the influence of cell motility and a target shape index were quantified. However, it should be noted that our model parameters cannot be directly mapped to this phase diagram which was constructed for confluent tissue represented with vertex models. Since these models assume a fully confluent tissue, adhesion and cortical tension are in direct competition, with adhesion favoring liquid-like behavior and cortical tension favoring jamming. In contrast, in our active particle simulations, adhesion and cortical tension both favor the glassy state, as a decrease in adhesion favors debonding and induces liquid-like tissue properties – SI Fig. 7.

On the aggregate level, the addition of Rho kinase inhibitor appears to cool down the tissue by reducing cell activity, hence inducing glassy behavior. Biologically, the inhibition of ROCK promotes actin depolymerization through cofilin, and reduces acto-myosin contractility by inhibition of myosin light chain phosphorylation [47]. These processes contribute to the active force generation mechanism of the cell, which might explain the role of Rocki as a modulator of the agitation level in a granular compaction model.

Y-27632 Rho kinase inhibitor is commonly used in low concentrations for bone and cartilage tissue engineering applications, where it is known to promote chondrogenic and osteogenic differentiation [48]. Among others, this effect has been attributed to biophysical factors, such as the inducement of cell rounding that leads to more porous aggregate structures facilitating mass transport [49]. This is consistent with our observation that ROCK inhibition produces aggregates that behave more like granular piles compared to a confluent fluid in control conditions. Mechanical stimulation and the presence of shear stress has been connected to the modulation of cell lineage commitment for control of tissue development following the initial mesenchymal condensation in various tissues such as bone [50] and tooth [51]. A key hallmark of the glassy state is the presence of long structural relaxation times. This entails that shear stress, which cannot be maintained in a fluid-like state, can be preserved at long times. The resulting mechanical environment, with the generation of cell-level shear stresses due to active cell forces, could mechanistically explain the necessity of actin inhibition during chondrogenic differentiation and its improved outcome in the presence of Rocki [52]. Taking these factors into account from the tissue engineering perspective, we propose that the design of in vitro tissues could be enhanced based on physical principles of granular matter in order to improve the robustness in outcomes that is currently hampering the field.

## Material and Methods

### Cell Expansion

Human periosteum-derived stem cells (hPDCs) were obtained from periosteal biopsies as described by Roberts et al. [53] The cells were expanded in T175 tissue culture flasks (5700 cells/cm^2^) in high glucose GlutaMAX™ Dulbecco’s modified Eagle’s medium (DMEM; Life Technologies, Merelbeke, Belgium) supplemented with 10% irradiated fetal bovine serum (FBS; HyClone), 1% sodium pyruvate (Invitrogen) and 1% antibiotic–antimycotic (100 units/ml penicillin, 100 mg/ml streptomycin, and 0.25 mg/ml amphotericin B; Invitrogen), defined as DMEM-c. Medium was changed every 2-3 days and cells were harvested with TrypLETM Express (Invitrogen) at 80%-90% confluency. During the whole process, the cells were incubated at 37 °C, 5% CO_2_ and 95% humidity inside a standard incubator. The ethical committee for Human Medical Research (KU Leuven) approved all procedures, and patients informed consent forms were obtained.

### Formation of micro-aggregates

A polydimethylsiloxaan (PDMS) micro-well mould (Dow Corning Sylgard 184 elastomer, MAVOM Chemical Solutions) was used for formation of micro-aggregates with homogenous size distribution. Micro-well inserts, containing micro-wells with a diameter of 200 µm and depth of 150 µm, were made by pouring 3% w/v agarose (Invitrogen, Belgium) onto the PDMS mould. The agarose was let to solidify where after 24-well inserts with an approximate area of 1.8 cm^2^ were punched out, containing approximately 2000 micro-wells. A detailed description of this protocol can be found in Leijten et al. [54]. Next, the inserts were fixed inside a 24-well plate with 1% w/v agarose and1 ml of DMEM-C was added to each well. The plate was sterilized under UV for 30 min and stored at 4 ^o^C until further use. For micro-aggregate production, cells were harvested and 100 or 200 cells/micro-aggregate were obtained by respectively seeding 1 ml of cell suspension at a concentration of 2×10^5^ or 4×10^5^ cells/mL. Micro-aggregates were cultured in DMEM-c with 0, 10 or 20 µM Rho-kinase inhibitor Y-276323 (Axon Medchem).

### Image acquisition and analysis

An integrated Live Cell Monitoring System (LiMSy), described in [6], was used to continuously monitor cell aggregation in an automated and high-throughput manner. Image acquisition was initialized by selecting two locations inside each well of interest. The focal point was manually configured for each frame and a time interval of 5 min was applied for an experiment that lasted 10 h in total. Next, the acquired data was analysed using the in-house developed software, composed of three major parts: (I) The micro-aggregates were automatically segmented from the images and their corresponding masks stored. (II) Features of interest, such as the area *A*_*a*_ and corresponding radius *R*_*a*_ = (*A*_*a*_*/π*)^1*/*2^, were extracted from the segmented masks. (III) Micro-well tracking was performed to monitor the aggregation behavior/response of single micro-aggregates. A more detailed description of the system and the software can be found in [6].

### Computational model

A detailed description of the individual cell-based computational model and the model setup can be found in the supplementary information.

## Acknowledgments

This work is part of Prometheus, the KU Leuven R&D Division for Skeletal Tissue Engineering. B.S. acknowledges support from the Research Foundation - Flanders (FWO), Grant Nr. 12Z6118N.

