## Supplementary Information for "Compaction dynamics during progenitor cell self-assembly reveal granular mechanics"

—


### Individual cell-based model

The individual cell-based computational model is loosely based on the concepts introduced in [1]. In this center-based model [2, 3], cells are represented by their central coordinates, and cell-cell contact interactions are computed for all edges of the Delaunay triangulation of these coordinates. This ensures that only nearest neighbours contacts may exist, even for densely packed tissue with soft repulsive potentials. For two cells $i$ and $j$ with positions $\boldsymbol{x}_i$ and $\boldsymbol{x}_j$ at a distance $d_{ij} = ||\boldsymbol{x}_j - \boldsymbol{x}_i||$ and with normal unit vector $\hat{\boldsymbol{n}}_{ij} = (\boldsymbol{x}_j - \boldsymbol{x}_i)/d_{ij}$, the central contact force reads:

$$\boldsymbol{F}^c_{ij} = \begin{cases} -w_{\mathrm{rep}}\, A_{c,ij}(d_{ij} - d^{\mathrm{eq}}_{ij})\,\hat{\boldsymbol{n}}_{ij} & \text{if } d_{ij} < d^{\mathrm{eq}}_{ij}, \\ -w_{\mathrm{adh}}\, A_{c,ij}(d_{ij} - d^{\mathrm{eq}}_{ij})\,\hat{\boldsymbol{n}}_{ij} & \text{if } d^{\mathrm{eq}}_{ij} \le d_{ij} \le d^{\max}_{ij}, \\ \mathbf{0} & \text{else} \end{cases} \tag{S1}$$

with $w_{\mathrm{rep}}$ the stiffness density effected by cortical tension and cell volume conservation and $w_{\mathrm{adh}}$ the attractive energy density effected by intercellar adhesion. $d^{\mathrm{eq}}_{ij} := c_{\mathrm{eq}}(R_i + R_j)$ is the contact's resting distance, where we use the ratio of the hexagon to the circle area to set $c_{\mathrm{eq}} = (\pi/(2\sqrt{3}))^{1/2}$, assuming that a 2D surface of equilibrated cells settles at a perfect hexagonal packing. $d^{\max}_{ij} := c_{\max}(R_i + R_j)$ describes the maximal range of adhesive interaction, and is influenced by the non-spherical shape of the cells, and its ability to form protrusions. In [1], $c_{\max}$ was empirically determined between 1.24 and 1.6, depending on the tissue type. We set $c_{\max} = 2^{1/3} \approx 1.26$, assuming that the surrounding volume where the cell can form interactions is equal to the cell's own volume. Active forces are also captured within contact interactions and are based on the cells' polarization directions $\hat{\boldsymbol{p}}_i$ and $\hat{\boldsymbol{p}}_j$. $A_{c,ij}$ represents the contact area between non-spherical fluid-like cells, and is computed using the quadratic expression $A_{c,ij} = a(d_{ij} - d_{ij}^{\max})^2$, with $a$ an empirical factor determined in [1] at $a \approx 1.3697$.

For a contact between $i$ and $j$, a protrusive force is exerted on cell $i$ if $\hat{\boldsymbol{n}}_{ij} \cdot \hat{\boldsymbol{p}}_i \geq \eta_{\min}$, a threshold value determining the width of the protrusive front. This force [1] reads:

$$\boldsymbol{F}_{ij}^{a} = T_a A_{c,ij} \left[ \left( \hat{\boldsymbol{p}}_i - \hat{\boldsymbol{p}}_j \right) \cos\nu + \left( \hat{\boldsymbol{p}}_i^{\perp j} - \hat{\boldsymbol{p}}_j^{\perp i} \right) \sin\nu \right], \tag{S2}$$

with

$$\hat{\boldsymbol{p}}_i^{\perp j} = \frac{\hat{\boldsymbol{n}}_{ij} - (\hat{\boldsymbol{n}}_{ij} \cdot \hat{\boldsymbol{p}}_i)\hat{\boldsymbol{p}}_i)}{||\hat{\boldsymbol{n}}_{ij} - (\hat{\boldsymbol{n}}_{ij} \cdot \hat{\boldsymbol{p}}_i)\hat{\boldsymbol{p}}_i||}.$$

$T_a$ is the maximal protrusive traction and $\nu$ is an angle that tunes the profile of the contact angle. In line with [1], we set $\eta_{min} = 0.15$ and $\nu = \arctan(4/3)$. To produce a persistent (angle) random walk, the polarization vector $\hat{\boldsymbol{p}}_i$ is perturbed each time increment $\Delta t$ by a rotation $\Delta\theta$ around a randomly chosen axis $\hat{\boldsymbol{u}}(t)$:

$$\Delta\theta_{i,\hat{\boldsymbol{u}}}(t) = \sqrt{2D_r\Delta t}\,\xi(t), \tag{S3}$$

where $\xi(t)$ is a typified Gaussian white noise. All force contributions are combined in the equation of motion for each cell as:

$$\sum_{j\in D(i)} \Gamma_{ij}(\dot{\boldsymbol{x}}_i - \dot{\boldsymbol{x}}_j) + \Gamma_{is}\dot{\boldsymbol{x}}_i + \gamma_{i,m}\dot{\boldsymbol{x}}_i = \boldsymbol{F}_i^{\mathrm{g}} + \boldsymbol{F}_i^{\mathrm{s}} + \sum_{j\in D(i)} (\boldsymbol{F}_{ij}^{c} + \boldsymbol{F}_{ij}^{a}), \tag{S4}$$

where $\gamma_{i,m} = 6\pi R_i \eta_{\mathrm{eff}}$ is the damping parameter from the effective viscosity of the medium $\eta_{\mathrm{eff}}$, and $\boldsymbol{F}_i^{\mathrm{g}} = V_i(\rho_{\mathrm{cell}} - \rho_{\mathrm{m}})\,\boldsymbol{g}$ is the net gravitational force due to the difference in cell $\rho_{\mathrm{cell}}$ and medium $\rho_{\mathrm{m}}$ density, and gravitational acceleration $\boldsymbol{g} = -g\,\hat{\boldsymbol{e}}_y$. $D(i)$ denotes all Delaunay neighbors (edges) of node $i$. $\boldsymbol{F}_i^{\mathrm{s}}$ is the cell-substrate contact force. Since adhesion between cells and agarose is assumed to be negligible, we model this as a simple Hertzian repulsion:

$$\boldsymbol{F}_i^{\mathrm{s}} = \begin{cases} \frac{4}{3}\hat{E}\,\hat{R}_{i,s}^{1/2}\,\delta_{i,s}^{3/2}\,\hat{\boldsymbol{n}}_w & \text{if}\,\delta_{i,s} > 0 \\ \boldsymbol{0} & \text{else,} \end{cases} \tag{S5}$$

with $\delta_{i,s}$ is the overlap distance with the substrate and $\hat{\boldsymbol{n}}_w$ is the outward normal unit vector to the microwell substrate. $\hat{E}$ is the effective stiffness of cell-agarose contact and $\hat{R}_{i,s}$ is the effective contact radius, which is $R_i$ for cell-bottom contact and $-R_iR_w/(R_i - R_w)$ for cell-wall contact, with $R_w$ the microwell radius. The cell-cell or cell-substrate resistance (friction) tensor is formulated as

$$\Gamma_{ij} = A_{c,ij}\left[\gamma_n \hat{\boldsymbol{n}}_{ij} \otimes \hat{\boldsymbol{n}}_{ij} + \gamma_t(\boldsymbol{I} - \hat{\boldsymbol{n}}_{ij} \otimes \hat{\boldsymbol{n}}_{ij})\right]. \tag{S6}$$

Here $\gamma_n$ and $\gamma_t$ represent normal (central) and tangential friction constants. For cells, these effective constants are the net result of the apparent cytosolic viscosity and the dynamic balance of intercellular adhesive ligands upon relative displacement of the cells. We do not investigate the relative contributions of $\gamma_n$ and $\gamma_t$ in detail. Rather we further assume that $\gamma_n = 2\,\gamma_t$, reasoning that the potentiality of intercellular sliding introduces an additional degree of freedom, causing $\gamma_n \geq \gamma_t$. For cell-substrate contacts, the contact area is computed according to Hertz' theory $A_{c,is} = \pi\delta_{i,s}\hat{R}_{i,s}$. (S4) can be written as a coupled system of equations:

$$\boldsymbol{\Gamma} \cdot \dot{\boldsymbol{x}} = \boldsymbol{F}. \tag{S7}$$

Here, $\boldsymbol{\Gamma}$ is a sparse positive definite resistance matrix with off-diagonal contributions due to cell-cell and cell-substrate contact interactions. Each time increment, this system is solved iteratively for the velocities using the conjugate gradient method. Time integration is performed in a semi-implicit scheme for first order systems. The cell positions are updated as

$$\boldsymbol{x}_{i,t+\Delta t} = \boldsymbol{x}_{i,t} + \Delta t\,\dot{\boldsymbol{x}}_{i,t+\Delta t}, \tag{S8}$$

where $\dot{\boldsymbol{x}}_{i,t+\Delta t}$ is the updated cell velocity that was obtained from solving the linear system (S7).

Experimentally, we compare simulation results to time-lapse microscopy data. This implies that we only have access to 'length' and 'time' data but not 'force'. To resolve this, we estimate relative (rescaled) parameters, relative to the cell-cell sliding viscosity $\gamma_t$. This leads to the following rescaled parameters:

$$\begin{aligned}\overline{\gamma}_n &:= \gamma_n/\gamma_t\\ v_t &:= T_a/\gamma_t\\ w_a &:= w_{\text{adh}}/(\overline{\gamma}_n\,\gamma_t)\\ w_r &:= w_{\text{rep}}/(\overline{\gamma}_n\,\gamma_t)\end{aligned}$$

Here, $v_t$ (μm/min) is the maximal instantaneous relative active velocity between two contacting cells, $w_a$ and $w_r$ (min$^{-1}$) are relaxation rates that determine how fast the distance between two contacting cells approaches its equilibrium value. Together with the rotational diffusivity $D_r$ (min$^{-1}$), $v_t$ and $w_a$ are the main influential parameters that determine the signature of the aggregate formation process. We further define the effective cell diffusivity as a net measure for cell activity (see Fig. 6 and 8): $D_{\text{eff}} := v_t^2/(2D_r)$.

### Simulation setup

The simulation of micro-well aggregation is split into two distinct phases: 1) a settling phase, during which the cells will sediment due to gravity on the bottom of the microwell and 2) the aggregation phase, during which the sedimented cells cohere and clump into a roughly spherical cell aggregate. We simulate both phases with the same individual cell-based model, albeit with some model parameters adjusted to the timescale of each process. During the short time (seconds or less) it takes for a cell to gravitationally settle at the bottom of the well, no adhesions have been formed yet, hence we set $w_\mathrm{a} = 0$ min$^{-1}$ and $v_t = 0$ μm/min. Consequently, the contact friction during this process does not represent the dynamic balance of bond breaking and creation, nor the viscous deformation of the cell, but simply the physical sliding of the non-adhering cells over each other. Therefore, we fix $\gamma_t = 0.02$ Pa·s/μm, 50 times lower than during the aggregation phase.

During the settling phase, cells are randomly generated at a fixed rate at the top of the micro-well. Assuming that in the experimental setup, the cells are uniformly distributed in the pipetted medium, the interval between the generation of two subsequent cells is on average $\Delta t_\mathrm{gen} = V_m/(A_m\, v_\mathrm{sed})$, with $V_m$ the total volume of pipetted suspension, $A_m$ the total area of the (macro-)well in which the suspension is pipetted and $v_\mathrm{sed}$ the average sedimentation velocity. The latter can be obtained from balancing Stokes' law to the gravitational force

$$v_\mathrm{sed} = \frac{g\,(\rho_c - \rho_m)V_c}{6\pi\eta_\mathrm{eff} R_c}. \tag{S9}$$

with $\rho_c$, $V_c$ and $R_c$ resp. the mass density, volume and radius of an average cell. To each generated cell $i$, we assign a radius $R_i$ randomly drawn from a list of measured cell radii (see Fig. 4), and a random position uniformly distributed in a disk with radius $R_w - R_i$. Apart from better approximating the experimental setting, the use of non-identical cell sizes has the additional benefit of preventing the emergence of artificial crystalline phases. This process is repeated until $N$ cells are added in the simulation. We further simulate for another 60 s to ensure that all cells are fully sedimented, and finalize the aggregation phase. A list of parameters used in the cell settling phase is given in Table 1.

Table 1: Simulation parameters that are unique to or specifically set for the simulation of the initial settling phase. Parameters that are not indicated in this table are identical to the ones further given in Table 2.

| **Parameter** | **Symbol** | **Value** | **Units** |
|---|---|---|---|
| Tangential friction | $\gamma_t$ | 0.02 | kPa·s/μm |
| Adhesive relaxation rate | $w_a$ | 0 | min$^{-1}$ |
| Relative active velocity | $v_t$ | 0 | μm/min |
| Viscosity medium | $\eta_\mathrm{eff}$ | 0.8 | mPa·s |
| Gravitational acceleration | $g$ | 9.81 | m/s$^2$ |
| Total medium area | $A_\mathrm{m}$ | 1.8 | cm$^2$ |
| Total medium volume | $V_\mathrm{m}$ | 1.0 | ml |
| Aggregation time | $t_a$ | 10 | h |
| Timestep | $\Delta t$ | 8 | ms |

Table 2: Simulation parameters used for the individual cell-based model, with indication of the source or rationale for each used value. 'Free' parameters that are studied in this work are not reported in this table.

| **Parameter** | **Symbol** | **Value** | **Units** | **Source** |
|---|---|---|---|---|
| Microwell radius | $R_\mathrm{w}$ | 108.56 | µm | Measured |
| Microwell height | $h_\mathrm{w}$ | 2000 | µm | Micro-well design |
| Average cell radius | $\langle R \rangle$ | 8.99 | µm | Measured |
| Standard deviation cell radius | $\sigma(R)$ | 1.42 | µm | Measured |
| Number of cells | $N$ | 200 | - | Set |
| Cell density | $\rho_c$ | 1100 | kg/m$^3$ | [4] |
| Medium density | $\rho_m$ | 1000 | kg/m$^3$ | |
| Effective medium viscosity | $\eta_\mathrm{eff}$ | 100 | mPa·s | |
| Gravitational acceleration | $g$ | 9.81 | m/s$^2$ | |
| Effective contact stiffness | $\hat{E}$ | 888 | Pa | Estimated |
| Tangential friction | $\gamma_t$ | 1.0 | kPa·s/µm | Reference value |
| Relative Normal friction | $\overline{\gamma}_n$ | 2.0 | $\gamma_t$ | Estimated |
| Cell-cell adhesion | $w_a$ | 4.0 | min$^{-1}$ | Estimated |
| Rotational diffusivity | $D_r$ | 0.5 | min$^{-1}$ | Estimated, see Fig. 6 |
| Attractive reach | $c_\mathrm{max}$ | $2^{1/3}$ | - | Estimated, [1] |
| Contact area factor | $a$ | 1.3697 | - | [1] |
| Protrusion tuning angle | $\eta_\mathrm{min}$ | 0.15 | - | [1] |
| Relative resting distance | $c_\mathrm{eq}$ | $(\pi/(2\sqrt{3}))^{1/2}$ | - | [1] |
| Aggregation time | $t_a$ | 10 | h | Set |
| Timestep | $\Delta t$ | 0.25 | s | Optimized |

Snapshots of the positions and radii of all cells are saved every 5 min time increment for each simulation. These are further processed to compute the aggregate area by drawing the cells as two-dimensional disks in the horizontal plane, and computing the intersection between these disks and the cells of a rectangular grid, where no single grid element can contain more than its own area. This procedure is visualized in Fig. 5. Hereafter, the temporal evolution of the aggregate area is obtained for each aggregate. This data is further analyzed in a manner that is identical to the analysis of experimental time-lapse microscopy data, to prevent artifacts due to differential data processing.

All simulations were performed in the particle-based simulation software Mpacts. For solving (S7), the conjugate gradient implementation of Eigen3 was used [5]. The initialization of the setup was performed in Python, and subsequent statistical data analysis was performed using the Numpy and Scipy packages in Python. All curve fitting procedures were carried out with the "curve_fit" subroutine that uses the Levenberg-Marquardt algorithm for nonlinear least square fitting. Visualizations of simulations were created in Paraview [6].

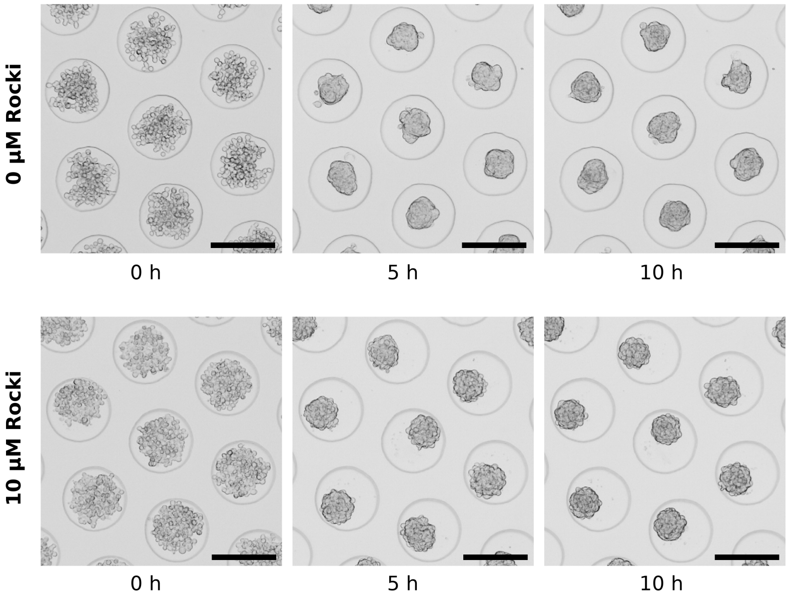


Figure 1: **Effect of Rho kinase inhibitor** Representative microscopy images of compacting aggregates of hPDCs at 0h, 5h, and 10h post-seeding, comparing the treatment with 0 and 10 µM of Rho kinase inhibitor (Rocki). The scale-bar is 200 µm.

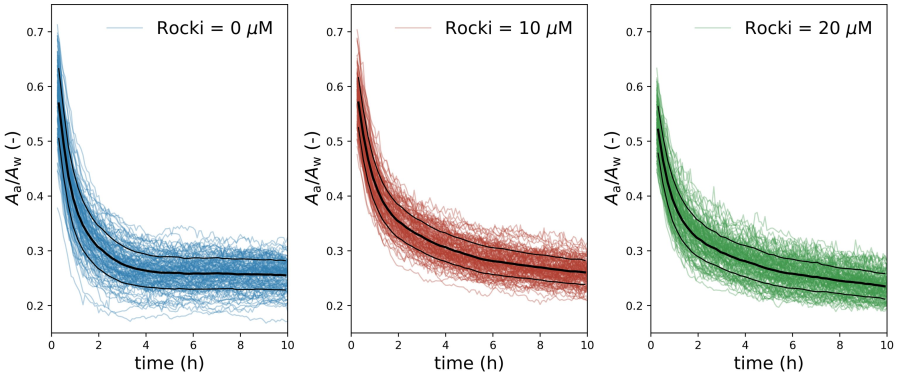


Figure 2: **Micro-aggregate area over time** for concentrations of Rho kinase inhibitor (Rocki) of 0 μM, 10 μM, and 20 μM, sampled every 5 minutes during the aggregate compaction process. Each colored line represents the area of a single analyzed micro-aggregate. The thick and thin black lines are resp. the average and ± the standard deviation over 195 (0 μM Rocki), 141 (10 μM Rocki) and 145 (10 μM Rocki) successful analyses.

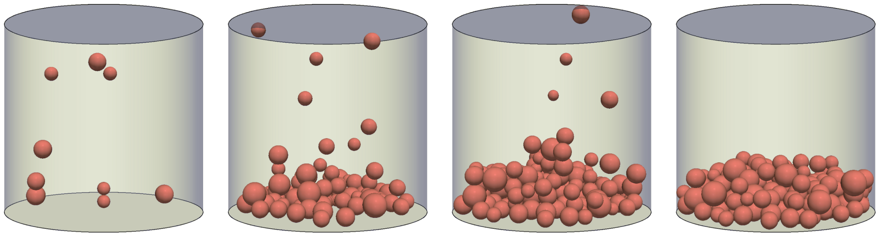

Figure 3: **Simulation of micro-well cell sedimentation.** Deposition simulation to describe the initial sedimentation which causes the initial structural configuration to start cell aggregation. Here, 200 cells are deposited in a microwell of approximately 200 μm diameter. From left to right: $t = 10\,\mathrm{s}$, $t = 110\,\mathrm{s}$, $t = 210\,\mathrm{s}$ and $t = 310\,\mathrm{s}$. Cell radii are sampled from measured cell sizes, as indicated in Fig. 4.

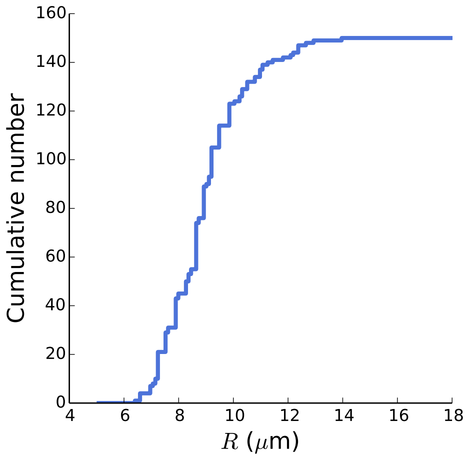


Figure 4: **Cumulative distribution of measured cell radii.** Distribution of 150 experimentally measured cell radii. The cell radii were assembled from microscopy images of progenitor cells seeded in microwells at low cell density, when isolated and highly circular cells can be easily distinguished. Cells radii were obtained using the 'circle' tool in image processing software imageJ.

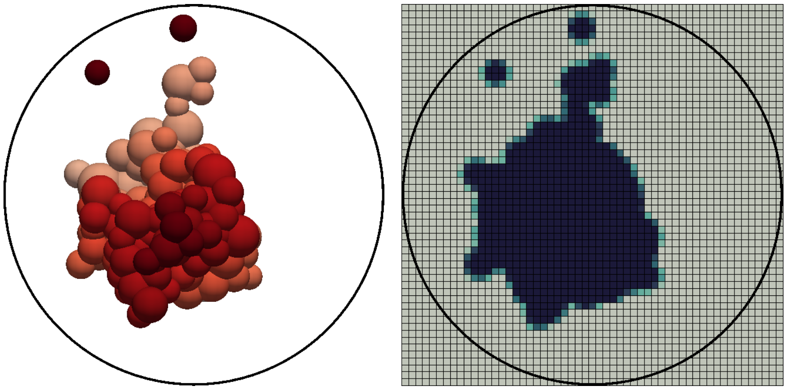

Figure 5: **Segmentation of aggregate area from simulations** Example of segmentation of a simulated aggregate (shown left). Each cell is drawn as a two-dimensional disk which is superposed on a rectangular grid of 4 µm × 4 µm, as shown on the right. The intersection between each disk and every grid cell (pixel) is computed and the resulting value is summed up to each pixel. The minimum of this sum and the pixel area is stored as the area value for each 'pixel'. The sum of all pixel values yields the estimated aggregate area. The large black circles indicates the micro-well with a radius of 100 µm.

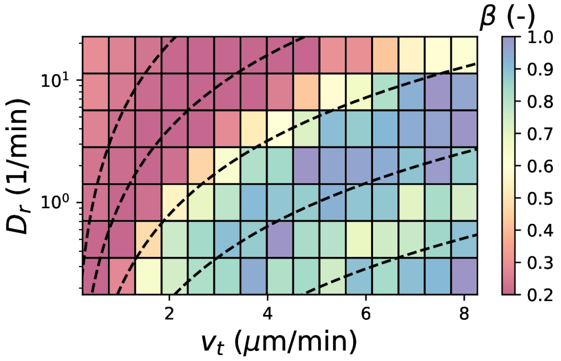


Figure 6: Simulated values of the exponent $\beta$ (see (2)) for varying active cell velocity $v_t$ and the rotational diffusivity of the cell $D_r$. Dashed guidelines show iso-contours of equal apparent cell diffusivity $D_{\text{eff}} = v_t^2/(2D_r)$, with values from left to right: 0.1, 0.5, 2.5, 12.5 and 62.5 μm$^2$/min. The trend in $\beta$ highly aligns with these iso-contours. This justifies the choice to consider $D_{\text{eff}}$ as a single measure for cell activity. Throughout the manuscript, we modulate $D_{\text{eff}}$ by varying $v_t$ and keeping $D_r$ constant at 0.5 min$^{-1}$. The latter value was chosen to represent a typical characteristic time of transient lammellipodia which last between 30 seconds and several minutes [7].

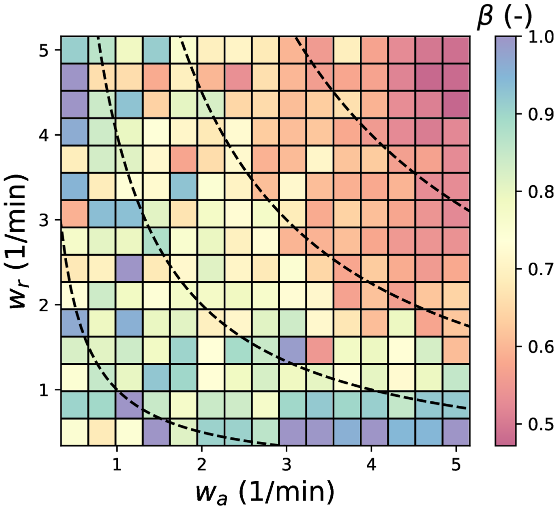


Figure 7: Map of estimated exponent $\beta$ of (2), for varying adhesion $w_a$ and repulsive stiffness $w_r$, simulated at $D_{\text{eff}} = 4$ μm$^2$/min and $D_r = 0.5$ min$^{-1}$. The dashed guidelines show iso-contours of $\sqrt{w_a w_r}$, the geometric mean of adhesive and repulsive stiffness, with values from bottom left to top corner: 1, 2, 3, and 4 min$^{-1}$.

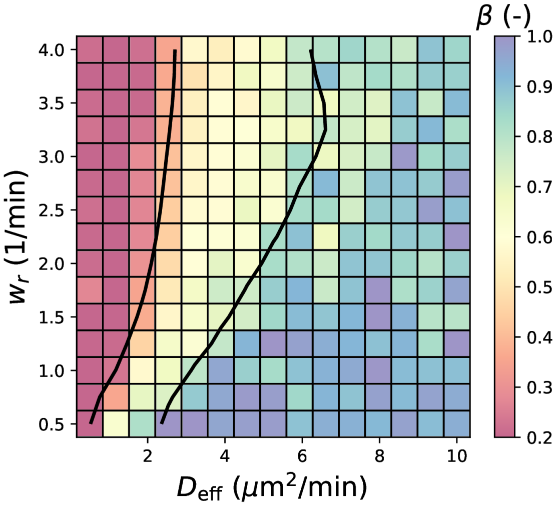


Figure 8: Map of estimated exponent $\beta$ of (2), for varying stiffness $w_r$ and cell activity $D_{\text{eff}} = v_t^2/(2D_r)$, simulated with a two times higher value of $D_r$ compared to Fig. 4(e), i.e. $D_r = 1\ \text{min}^{-1}$, but keeping all other parameters identical.

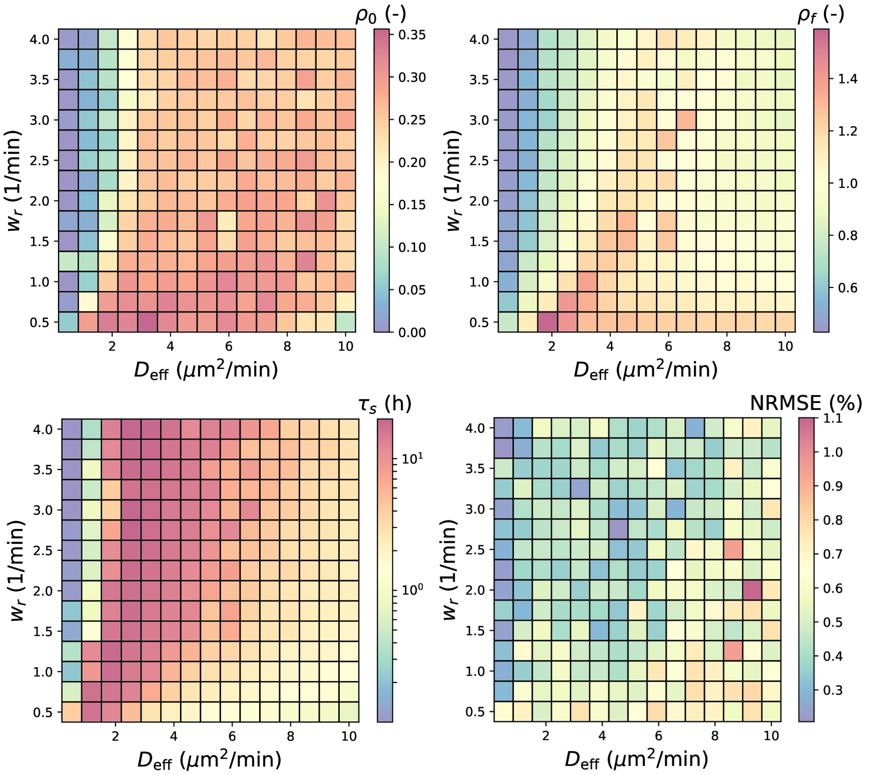


Figure 9: Maps of additional fitted parameters $\rho_0$, $\rho_f$, $\tau_s$ from (2) for varying stiffness $w_r$ and cell activity $D_{\text{eff}} = v_t^2/(2D_r)$, corresponding to the simulation settings used for Fig. 4(e). The NRMSE (normalized root mean square error, bottom right) indicates that (2) provides a good fit for the dynamics of compaction for all simulated parameter combinations.
